# System-level analysis of genes mutated in muscular dystrophies reveals a functional pattern associated with muscle weakness distribution

**DOI:** 10.1101/2024.01.05.574331

**Authors:** Ozan Ozisik, Svetlana Gorokhova, Mathieu Cerino, Marc Bartoli, Anaïs Baudot

## Abstract

Muscular dystrophies (MDs) are inherited genetic diseases causing weakness and degeneration of muscles. The distribution of muscle weakness differs between MDs, involving distal muscles or proximal muscles. While the mutations in most of the MD-associated genes lead to either distal or proximal onset, there are also genes whose mutations can cause both types of onsets.

We hypothesized that the genes associated with different MD onsets code proteins with distinct cellular functions. To investigate this, we collected the MD-associated genes and assigned them to three onset groups: genes mutated only in distal onset dystrophies, genes mutated only in proximal onset dystrophies, and genes mutated in both types of onsets. We then systematically evaluated the cellular functions of these gene sets with computational strategies based on functional enrichment analysis and biological network analysis.

Our analyses demonstrate that genes mutated in either distal or proximal onset MDs code proteins linked with two distinct sets of cellular processes. Interestingly, these two sets of cellular processes are relevant for the genes that are associated with both onsets. Moreover, the genes associated with both onsets display high centrality and connectivity in the network of muscular dystrophy genes. Our findings support the hypothesis that the proteins associated with distal or proximal onsets have distinct functional characteristics, whereas the proteins associated with both onsets are multifunctional.

## INTRODUCTION

Muscular dystrophies (MDs) are inherited genetic diseases characterized by the weakness and degeneration of muscles [1]. They form a highly heterogeneous group of diseases, with great variations in age of onset, severity and progression. The groups of muscles primarily affected in each MD as well as associated non-muscular phenotypes are also highly heterogeneous [2]. These different features are used for the clinical classification of MDs. This classification considers for instance congenital or later onset conditions, but also the localization and distribution of muscle weaknesses. Indeed, muscle weakness distribution is an important feature of MDs [3]. In particular, distal and proximal onsets are observed in MD patients. In distal onset, muscle weakness is primarily observed in the distal parts of the upper and lower limbs (hands and feet); whereas, in proximal onset, muscle weakness is primarily observed in the muscles that connect the limbs to the trunk.

At least 40 genes are currently known to cause MDs when mutated [1]. The identification of mutations and mutated genes helped to refine the classification of MDs and investigate their underlying molecular pathomechanisms. The genes mutated in MDs encode proteins involved in a wide variety of muscle-related functions (e.g. muscle development, contractility, elasticity) and cellular localizations (e.g. extracellular matrix, nuclear membrane) [1, 3]. A recent review article suggested that the biological processes and cellular localizations of the proteins involved in MD pathogenesis would determine the muscle weakness distribution, the distal and proximal onsets in particular [3]. More specifically, it was hypothesized that mutated proteins located in sarcomere and Z-disk would cause distal weakness, whereas abnormal proteins located in muscle cell membrane (sarcolemma, sarcoplasmic reticulum, nuclear membrane) would cause proximal weakness [3]. For example, MYH7 protein is located in sarcomere and pathogenic variants in *MYH7* gene are responsible for Distal Myopathy 1 (MPD1, also known as Laing distal myopathy), whereas, A-type lamins are located at the inner nuclear membrane and pathogenic variants in *LMNA* gene are responsible for Emery-Dreifuss muscular dystrophy2 (EDMD2) that has a proximal onset. Interestingly, some genes can cause both proximal and distal onset MDs when mutated, e.g. *DES, DYSF, MYOT, TTN*. Multifunction and/or dual localization has been suggested as an explanation to this phenomenon [3].

To our knowledge, no study has performed a systematic, unbiased analysis of the functional characteristics of the genes associated with different MD onsets. In this study, we conducted this analysis using two systems biology strategies: functional enrichment analysis and network analysis.

Functional enrichment analysis identifies shared functions among a given set of genes or proteins by incorporating the biological knowledge represented as predefined annotations, such as Gene Ontology (GO) terms [4, 5]. Network analysis, on the other hand, goes beyond these predefined annotations. Using the molecular interaction information, it can assess whether two proteins have a shared role in a biological process or a condition. The underlying rationale for this is that proteins participating in the same biological processes cluster in close network vicinity [6]. Consequently, mutated genes coding for interacting proteins might lead to the same or similar phenotypes [6, 7, 8].

Here, based on systematic enrichment and network analyses, we investigated the common and distinct functional characteristics associated with different MD onsets. We revealed that proteins associated with either distal or proximal onsets have distinct functional characteristics, whereas the proteins associated with both distal and proximal onsets are multifunctional.

## MATERIALS AND METHODS

### Collection of genes associated with distal and proximal onset muscular dystrophies

French National Network for Rare Neuromuscular Diseases (*Filière Nationale des Maladies Rares Neuromusculaires*, FILNEMUS) has previously established lists of genes to be analyzed based on the clinical diagnosis entry groups [9]. We extracted the genes from the following two lists: “Distal and Scapuloperoneal Myopathies - exhaustive list” and “Limb Girdle Muscular Dystrophies - exhaustive list”. These two lists contain genes associated with distal and proximal onset muscular dystrophies (MDs), respectively. Krahn et al. estimated the Gene-Disease associations as “Convincing” or “Limited” [9]. However, these associations were established before the extensive curation efforts became available [10, 11, 12, 13]. We therefore also queried The GenCC database [13] (https://thegencc.org, accessed on 13.02.2023). We retained the genes from the FILNEMUS lists only if they were curated in the GenCC database as “Moderate/Strong/Definitive” and for an MD with the same onset as indicated in FILNEMUS. The following genes were also retained given the extensive literature support and “Convincing” curation by Krahn et al., even though they have not yet been curated as “Moderate/Strong/Definitive” in the GenCC database: *CRYAB, MATR3, MYOT*.

Finally, we assigned these 40 MD-associated genes to three onset groups: Distal-only Genes (DoGs) that contain the genes exclusively associated with distal onset MDs (present only in the “Distal and Scapuloperoneal Myopathies” list); Proximal-only Genes (PoGs) that contain the genes exclusively associated with proximal onset MDs (present only in the “Limb Girdle Muscular Dystrophies” list); and Common Genes (CoGs) that contain the genes that are associated with both types of onsets (present on both lists) (Table 1).

**Table 1.**
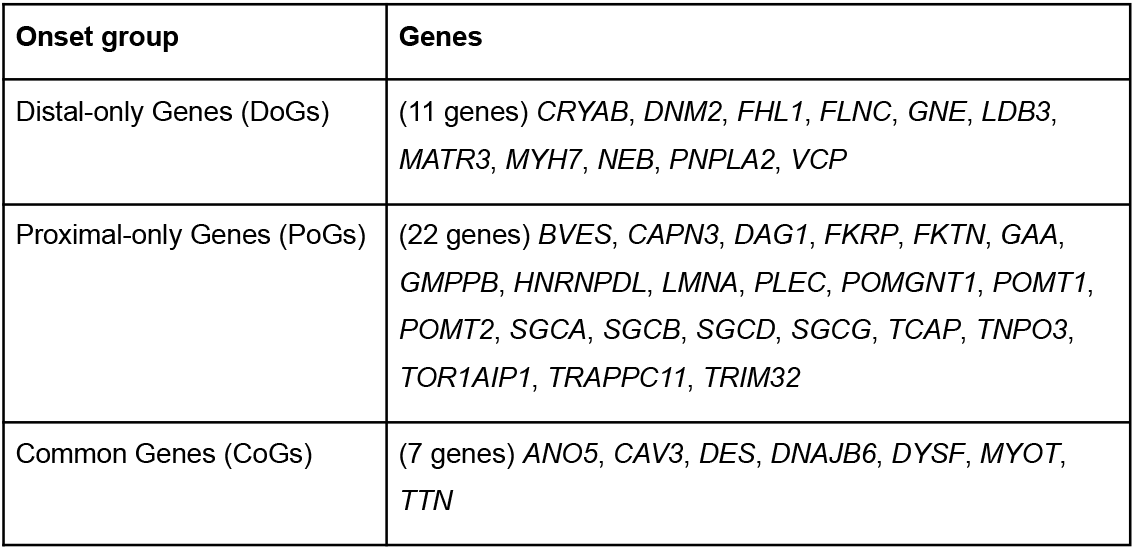
Muscular dystrophy onset groups and associated genes.

| Onset group | Genes |
| --- | --- |
| Distal-only Genes (DoGs) | (11 genes) *CRYAB*, *DNM2*, *FHL1*, *FLNC*, *GNE*, *LDB3*, *MATR3*, *MYH7*, *NEB*, *PNPLA2*, *VCP* |
| Proximal-only Genes (PoGs) | (22 genes) *BVES*, *CAPN3*, *DAG1*, *FKRP*, *FKTN*, *GAA*, *GMPPB*, *HNRNPDL*, *LMNA*, *PLEC*, *POMGNT1*, *POMT1*, *POMT2*, *SGCA*, *SGCB*, *SGCD*, *SGCG*, *TCAP*, *TNPO3*, *TOR1AIP1*, *TRAPPC11*, *TRIM32* |
| Common Genes (CoGs) | (7 genes) *ANO5*, *CAV3*, *DES*, *DNAJB6*, *DYSF*, *MYOT*, *TTN* |

### Functional enrichment analysis

We performed functional enrichment analyses using the g:Profiler Python client (package version 1.0.0, g:Profiler version e109_eg56_p17_1d3191d, database updated on 29/03/2023) [14]. We used g:Profiler’s default parameters (statistical domain scope: only annotated genes; significance threshold: g:SCS threshold; user threshold: 0.05). We evaluated DoGs, CoGs and PoGs separately for their enrichments in Gene Ontology (GO) terms [4, 5], using biological process (GO:BP) and cellular component (GO:CC) domains. We filtered and compared the enrichment analysis results using the orsum Python package (version 1.6) [15]. orsum works with annotation databases that are hierarchically organized such that terms are in a parent-child relationship, e.g. GO. orsum selects more significant general terms as representatives for their less significant child terms. orsum also allows integrating results obtained in multiple independent analyses and cluster the results for easy interpretation. We ran orsum with the default parameters except that we set the number of annotation terms to be displayed in the plots to 20.

### Biological network data

Multiplex networks are networks composed of multiple layers sharing the same set of nodes but different interactions. We used a multiplex network composed of three layers of interactions: a protein-protein interaction (PPI) layer, a molecular complexes layer and a pathways layer. The PPI layer gathers binary physical interactions between proteins. It was built using 3 datasets: APID [16, 17] (Level 2, human only), Hi-Union and Lit-BM [18]. The molecular complexes layer gathers the interactions between the members of macromolecular machineries, identified for instance with purification experiments. It was built from the fusion of Hu.map [19] and CORUM [20]. The pathways layer was built using the Reactome protein-protein interaction data [21]. The networks were built following the procedures available at https://github.com/CecileBeust/Networks_building. The largest connected component of the multiplex network was extracted and used in all the analyses (see supplementary Table S1 and S2 for network metrics). It should be noted that one CoG (*ANO5*) and two PoGs (*FKRP* and *FKTN*) do not have any interaction in this multiplex network and were thereby not considered in the network analyses.

In order to verify the results obtained using the multiplex network with a different, single layer network, we used the BioGRID PPI network (version 4.4.222) [22]. In this network we kept only the physical interactions between human proteins.

We used Cytoscape 3.9.1 [23] for network visualization. We used NetworkAnalyzer in Cytoscape to calculate the network metrics such as network diameter and node centralities.

### Average shortest distances between onset groups

We computed the average shortest distances between genes belonging to the same onset group, i.e. DoGs, CoGs and PoGs (intra-group distances), as well as between the genes that belong to different onset groups (inter-group distances). Additionally, we created random gene sets whose sizes are similar to onset groups and calculated the average distances from the different onset groups to these random gene sets. We assessed whether the distances calculated for different onset groups are smaller than expected using a bootstrap approach: Let’s say we are assessing the distance from DoGs to PoGs; we create 2000 random gene sets containing the same number of genes as PoGs, calculate the distance of DoGs to these random sets, and compute the ratio of random gene sets that have a distance smaller than the real distance between DoGs and PoGs, just by chance. We use this ratio as the significance value of the distance from DoGs to PoGs. Note that this significance assessment is not symmetric; we indeed performed another assessment for the distance from PoGs to DoGs.

We used in-house Python code and the NetworkX Python package [24] in these analyses.

### Extraction of network neighborhoods by random walk with restart

Random walk with restart (RWR) calculates the probability that a walker, starting from one of the seed nodes, walking randomly on the network edges, and randomly restarting the walk from one of the seed nodes, will visit a specific node in the network. With RWR, each node in the network will obtain a probability, called RWR score from now on, that is based on both the node’s distance to the seed nodes and the topology of the network. The RWR algorithm was recently extended to explore multiplex and multilayer networks [25, 26].

We used the RWR scores to extract the network neighborhoods of the different onset groups. To this goal, we used the DoGs, CoGs and PoGs as seeds in three separate RWR runs. In each run, we collected the top-scoring 50 genes as the network neighborhood of the respective onset group. We additionally selected the top-scoring 100 genes to check whether the results are consistent with the results obtained when the top 50 genes are selected. To run the RWR, we used the MultiXrank Python package [26] with the default restart probability (0.7).

We checked the overlap of the network neighborhoods obtained by using different onset groups as seeds. Additionally, we assessed the likeliness of the overlap occurring by chance using a bootstrap approach: We randomized the labels (“DoG”, “CoG”, “PoG”) of the MD-associated genes 1000 times and compared the overlap obtained in each run with the overlap obtained in the real case. Finally, we reproduced the same analysis using a single layer network, BioGRID PPI network.

## RESULTS

### 40 genes are associated with distal and/or proximal onset muscular dystrophies

We obtained 40 genes associated with distal and proximal onsets from the study by Krahn et al. [9] and the GenCC database (Materials and Methods). We will refer to these genes collectively as muscular dystrophy-associated (MD-associated) genes. We next categorized these genes into three onset groups, namely Distal-only Genes (DoGs), Proximal-only Genes (PoGs) and Common Genes (CoGs), the last corresponding to the genes associated with both distal and proximal onset MDs (Table 1).

### Genes mutated in distal and proximal muscular dystrophies are associated with different functional enrichments

We performed functional enrichment analyses of the gene sets belonging to different onset groups, i.e. DoGs, CoGs and PoGs (Materials and Methods). At the level of biological processes (GO:BP), all the onset groups are enriched in “muscle structure development” (Figure 1). We can also observe enrichments specific to each onset group. In particular, DoGs and PoGs present distinct enrichment profiles. DoGs are enriched in “striated muscle cell development” and “actomyosin structure organization”. PoGs are enriched in processes related to cardiac development, mannosylation and glycosylation. Finally, CoGs are the only onset group significantly enriched in “muscle contraction” and “plasma membrane repair” (Figure 1).

**Figure 1.**
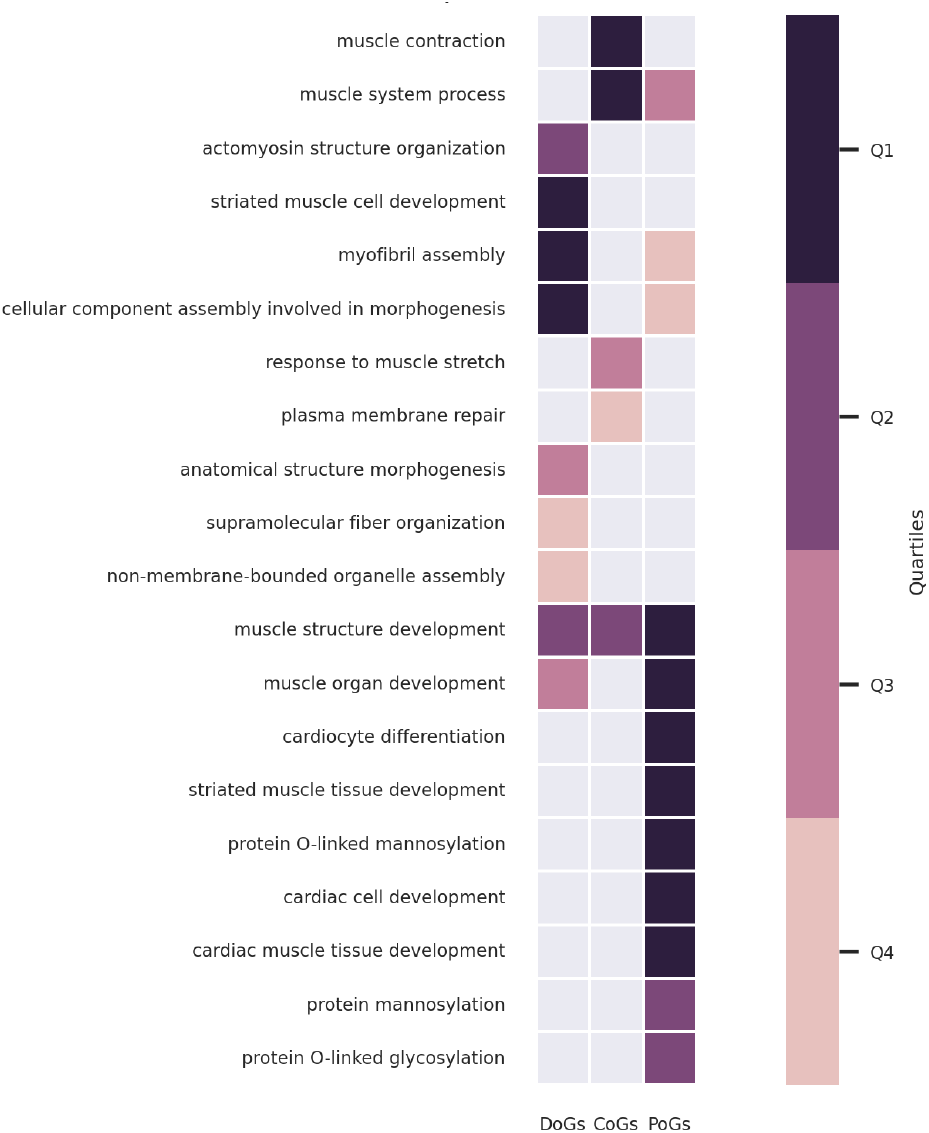
GO:BP enrichment analysis results for DoGs, CoGs and PoGs. The results are filtered and clustered using orsum. The colors represent the quartiles of the terms’ enrichment significance (darker means more significant).

At the level of cellular localization (GO:CC), all onset groups are enriched in “sarcomere”, “myofibril” and “contractile fiber” (Figure 2). However, it should be noted that PoGs are less significantly enriched in these terms as compared to DoGs and CoGs (Figure 2, supplementary enrichment results). DoGs and CoGs are also enriched in “Z disc” and “I band” cellular localizations. The most significant GO:CC terms for PoGs are “sarcolemma” and “glycoprotein complex” related terms. “Sarcolemma” is also significant for CoGs. The different enrichments in sarcolemma and sarcomere annotations in the different onset groups looked particularly interesting. Focusing only on the enrichments with these two annotation terms, we indeed observed very significant enrichments (adjusted p-value < 10^−5^) for DoGs in sarcomere and PoGs in sarcolemma localization. Contrarily, CoGs display very significant enrichments (adjusted p-value < 10^−5^) for both localizations (Table 2).

**Table 2.**
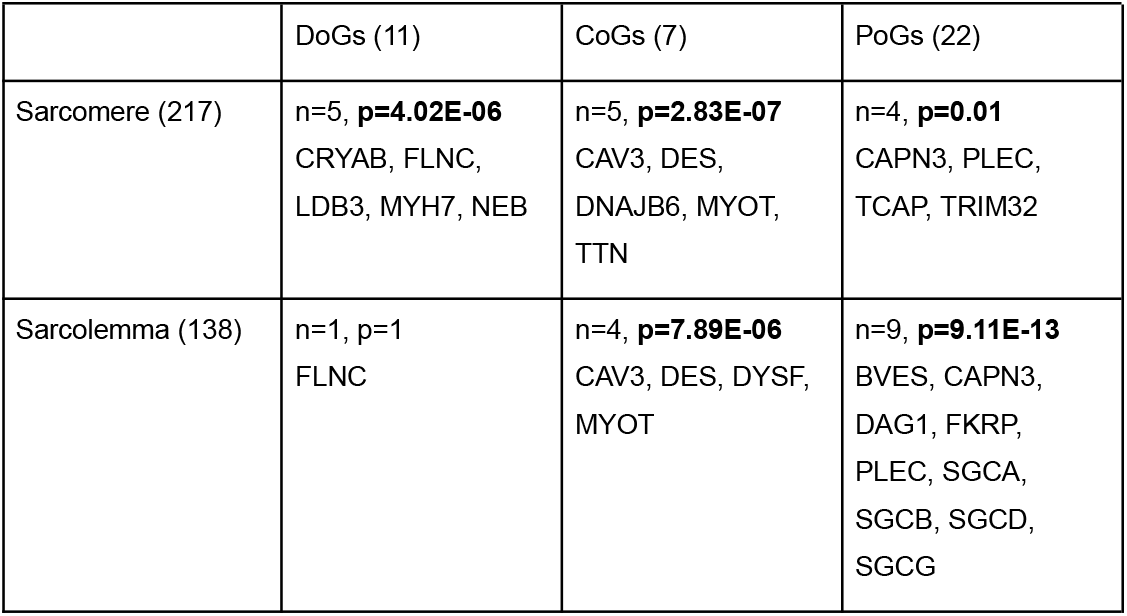
Enrichment statistics of DoGs, CoGs and PoGs in sarcomere and sarcolemma cellular localizations (GO:CC). The sizes of the gene sets are given in parentheses. The number of genes annotated for the given cellular localization (n) and the adjusted p-value of the enrichments (p) are indicated. Significant p-values are shown in bold.

**Figure 2.**
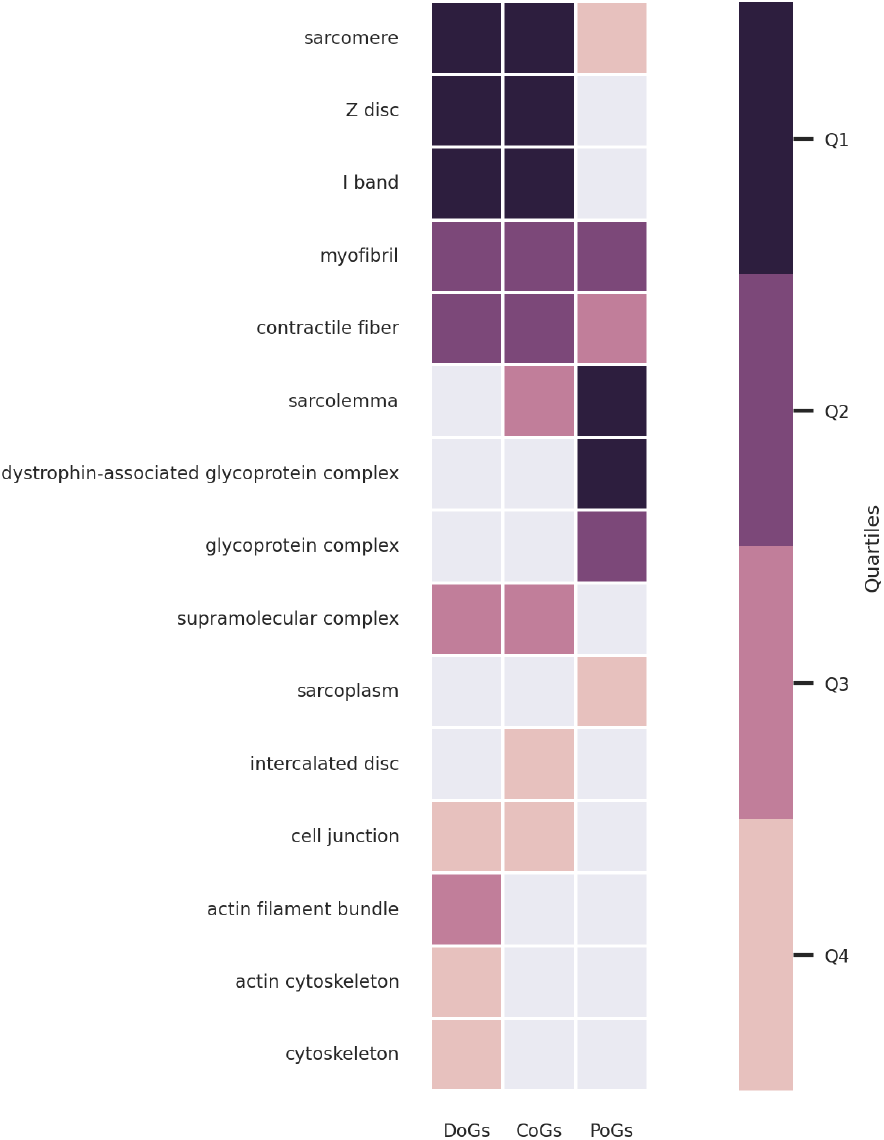
GO:CC enrichment analysis results for DoGs, CoGs and PoGs. The results are filtered and clustered using orsum. The colors represent the quartiles of the terms’ enrichment significance (darker means more significant).

These enrichment analyses overall reveal that genes mutated in distal and proximal onset MDs are associated with distinct biological processes and cellular localizations. Importantly, this suggests that, when mutated, sarcomeric proteins cause distal weakness and sarcolemmic proteins cause proximal weakness.

### CoGs are central in the subnetwork of muscular dystrophy-associated genes

We collected and visualized the direct interactions between the MD-associated genes (Figure 3). In the large connected component of the network presented in Figure 3, *DYSF* has the highest closeness centrality, and four of the five nodes with the highest closeness centrality are CoGs (supplementary Cytoscape session file Networks.cys). CoGs are hence highly central among MD-associated genes.

**Figure 3.**
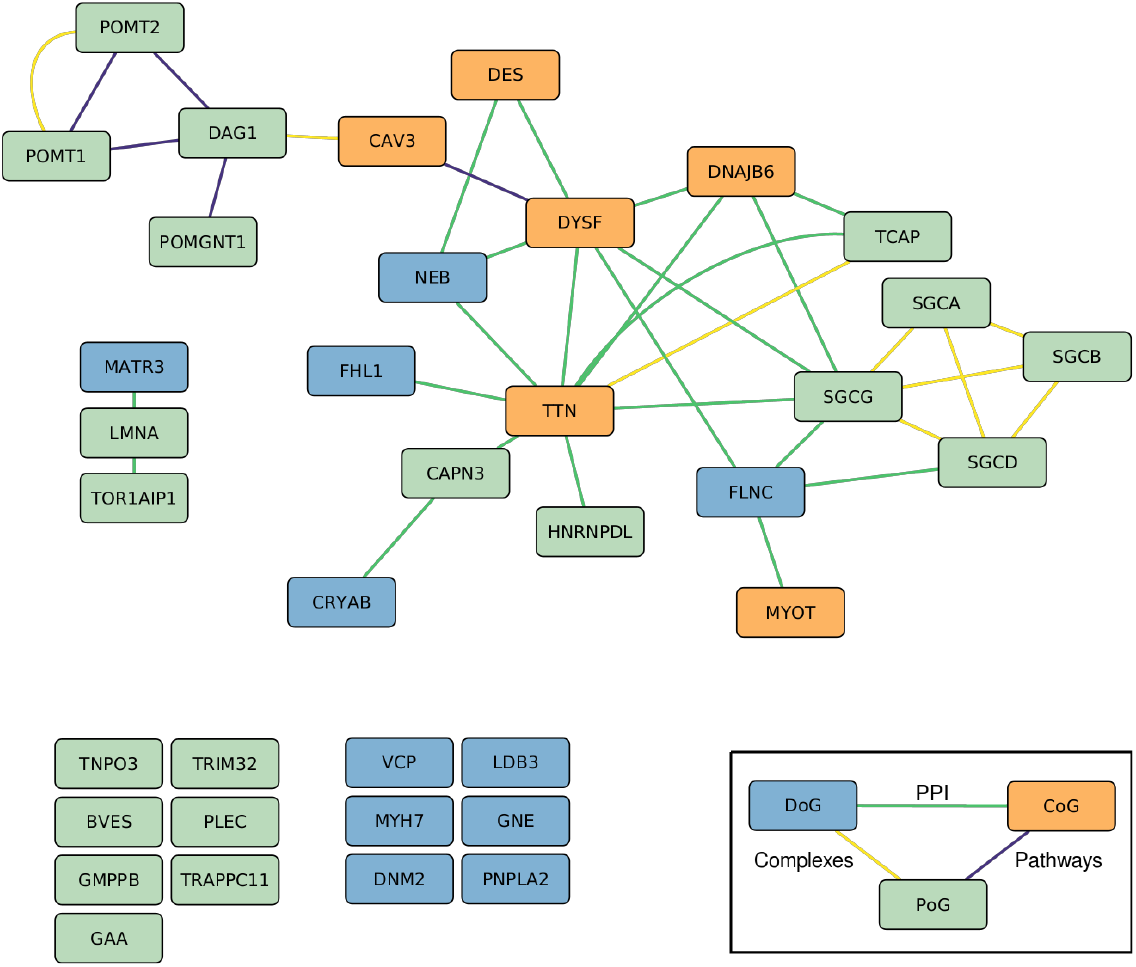
Direct interactions between MD-associated genes. For the sake of visualization, the different layers of the multiplex network (protein-protein interaction (PPI), Complexes, Pathways) are merged in a single layer. The disconnected nodes are MD-associated genes that do not have any direct interactions with other MD-associated genes, but they have other interactions within the multiplex network. Node colors represent the onset group; Distal genes (DoG) are shown in blue, Common genes (CoG) in orange and Proximal Genes (PoG) in green. Edge colors represent the interaction layer; PPI in green, complexes in yellow and pathways in violet.

### CoGs are central among muscular dystrophy-associated genes in the whole network

We next calculated the average shortest network distances between the genes belonging to the same onset group (intra-group distance), as well as in between the genes belonging to different groups (inter-group distance). In this case, the distances were computed on the full multiplex network (Materials and Methods). CoGs have the smallest intra-group distance, with an average shortest distance of 1.8 (Table 3). Interestingly, DoGs and PoGs also have shorter average distances with CoGs than with genes of their own onset group. These results reinforce the observation that CoGs are central among MD-associated genes in the network.

**Table 3.** Intra-group distances, inter-group distances and distances to random gene sets.

|  | DoGs | CoGs | PoGs | Random |
| --- | --- | --- | --- | --- |
| DoGs | 2.51 | 2.29 | 2.77 | 3.02 |
| CoGs |  | 1.8 | 2.51 | 3.04 |
| PoGs |  |  | 2.91 | 3.21 |
| Random |  |  |  | 3.42 |

We additionally created random sets of genes and calculated the average distances of onset groups to these random gene sets. The intra- and inter-group distances of all the onset groups are smaller than the average distances to random gene sets (Table 3). Finally, we assessed whether the calculated distances are smaller than expected using a bootstrap approach (Materials and Methods), and observed that they are significantly smaller than expected by chance (p ≤ 0.001). These results demonstrate that MD-associated genes are not scattered around the network, but rather localized close to each other.

### There is little overlap between the network neighborhoods of DoGs and PoGs

Nodes that are in close relation with MD-associated genes in the network are interesting because they might be affected the most when MD-associated genes are mutated. We used random walk with restart (RWR) to extract the network neighborhood of DoGs, CoGs and PoGs (Figure 4, supplementary network neighborhood results, Materials and Methods).

**Figure 4.**
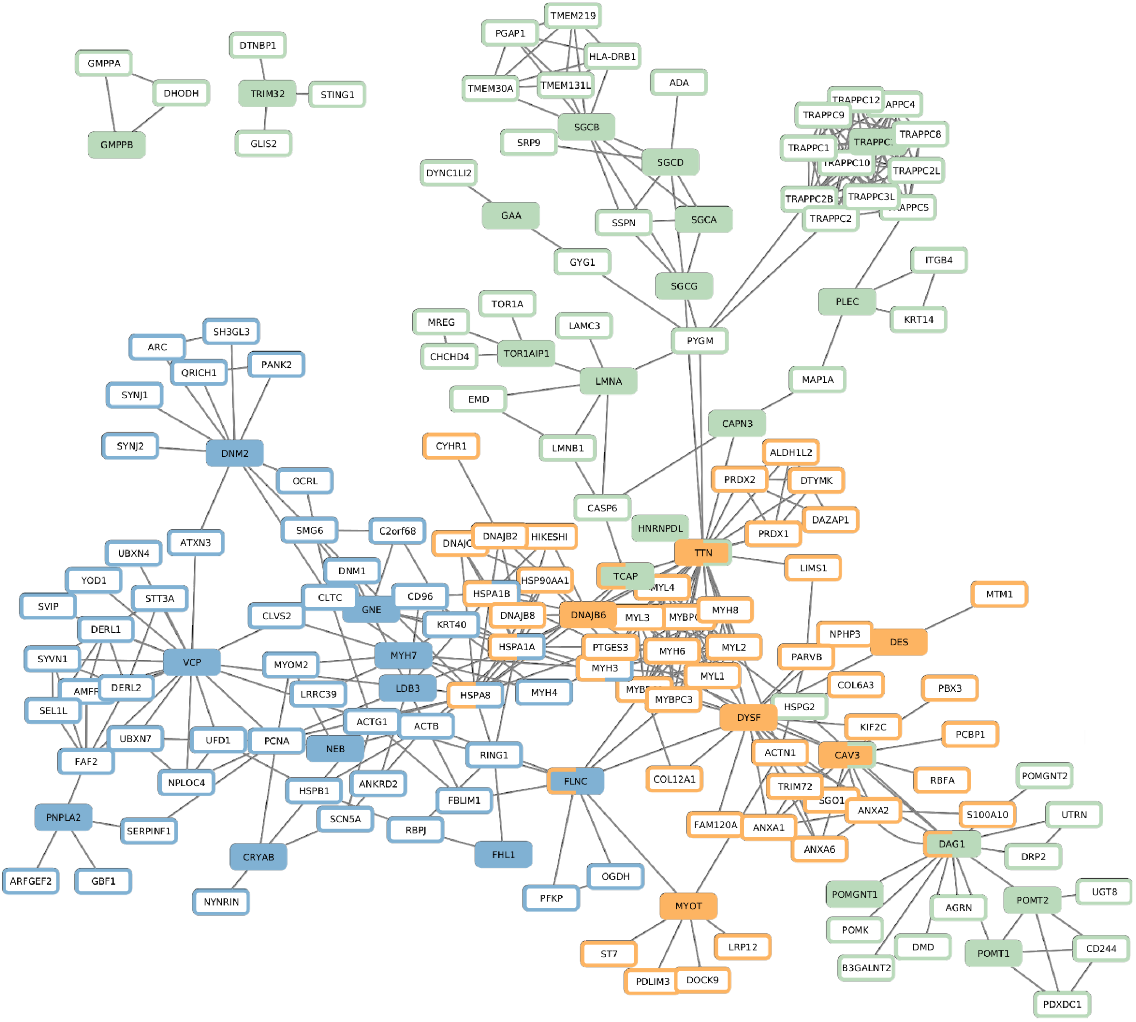
Network neighborhoods of DoGs, CoGs and PoGs (computed using the top 50 RWR scores on multiplex network). Filled nodes correspond to seeds, with colors indicating the onset groups (DoGs: blue, CoGs: orange, PoGs: green, as given in the legend of Figure 3). Node border colors indicate whether a node belongs to the neighborhood of DoGs, CoGs or PoGs. In the cases that seeds or neighborhoods of different onset groups overlap, both onset groups are indicated in the node borders.

We observed that the neighborhoods of DoGs and PoGs do not share any genes. On the other hand, the neighborhood of CoGs contains a DoG and two PoGs, and it overlaps with the neighborhood of DoGs.

In order to assess the significance of non-overlapping network neighborhoods of DoGs and PoGs, we randomized the gene labels (“DoG” and “PoG”), ran RWR, and checked the overlap of the neighborhoods. In 1000 randomizations, the ratio of overlaps less than or equal to the real overlap is 0.031.

In order to see if the results are consistent when different settings are used, we made two additional trials. First, we selected the top scoring 100 genes as the neighborhood, instead of the top 50. Second, we selected the top 50 genes as we did in our main analysis but we ran RWR on another biological network, the BioGRID PPI (Materials and Methods). In both settings, we observed some overlap between the neighborhoods of DoGs and PoGs but the overlaps were still less than what we would obtain by chance; in 1000 randomizations, the ratio of overlaps less than or equal to the real overlap was 0.043 for the first trial and 0.121 for the second trial.

These results demonstrate that there is little overlap between the network neighborhoods of DoGs and PoGs and, when mutated, they might affect different biological processes.

## DISCUSSION

Muscular dystrophies (MDs) encompass diverse genetic diseases characterized by different onsets of muscle weakness and degeneration. The muscle weakness distribution in MDs can differ, involving distal muscles, proximal muscles, or both. The differences in the physiology and the susceptibility to dysfunction of proximal vs. distal muscle groups are not clearly understood. MD-associated genes, regardless of the associated onset, are expressed in all skeletal muscles. Therefore, the differences in onset location cannot be explained by muscle-group specific gene expression. Instead, it is likely that distinct onset locations reflect the increased initial susceptibility of proximal or distal muscle groups to the disruption of certain biological processes.

We conducted a system-level analysis with two complementary approaches to investigate the functional characteristics of proteins associated with different MD onsets. We observed that genes mutated only in distal onset MDs (Distal-only Genes, DoGs) and genes mutated only in proximal onset MDs (Proximal-only Genes, PoGs) are associated with distinct cellular functions, characterized both from a functional enrichment perspective and a network analysis perspective. In addition, genes mutated in both distal and proximal onset MDs (Common Genes, CoGs) share functional characteristics with both of the other onset groups, i.e. DoGs and PoGs.

DoGs are enriched in terms related to sarcomere, “Z disc” being the most significant term, followed by “I band” and “sarcomere”. In contrast, PoGs are enriched in terms related to sarcolemma, in both cellular localization and function (e.g. “protein O-linked mannosylation”). CoGs, on the other hand, are enriched in both “sarcomere” and “sarcolemma” terms. These findings support the previously proposed hypothesis that dysfunctions of membrane-bound proteins cause proximal MDs, while intracellular proteins are more linked to the distal onset [3]. However, certain genes do not completely fit with this hypothesis, as they are associated with only one of the onsets but annotated with both “sarcomere” and “sarcolemma” terms (e.g. *FLNC, CAPN3, PLEC*). Moreover, two PoGs, *TCAP* and *TRIM32*, are annotated with the “sarcomere” term. One of the possible reasons for these results is that phenotypic data are still incomplete for recently described or very rare forms of MDs. For example, initially described cases with biallelic *TCAP* variants had proximal presentations, while more recent reports also describe patients with distal onset [27]. Genes might also have different splicing isoforms that have different functions in distinct subcellular localizations.

Our network analysis results are concordant with the enrichment analysis results. We observed that the network neighborhoods of DoGs and PoGs do not overlap, which indicates distinct functions. Another observation is that CoGs are highly central and well-connected among MD-associated genes. This suggests that CoGs may play a crucial role in mediating interactions among MD-associated genes. Pathogenic variants in CoGs are thus likely to perturb the processes that involve either DoGs or PoGs, thus leading to abnormal function spanning multiple muscle groups. Our observations that CoGs share functional enrichments with the two other onset groups is concordant with this.

We identified here different functional characteristics for the genes involved in MDs with different onsets. However, it is to note that other types of mechanisms could be involved in addition or in parallel. One possibility is that fiber type composition between muscle groups could underlie the differences in susceptibility to muscle wasting in various pathological conditions [28]. Indeed, muscles are composed of several types of muscle fibers that have different distribution patterns between muscle groups and within individual muscles. For example, the density of “slow” anaerobic type I fibers decrease from proximal toward more distal levels [29]. Moreover, in certain genetic muscle diseases one muscle type is preferentially affected [30]. Other proposed differences between proximal and distal muscles include variations in developmental processes and innervation as well as differences in thermal balance and pressure increase linked to muscle activity [29, 31]. Future research should further explore the fiber type compositions of different muscle groups, expression patterns of MD-associated genes in these fiber types, and also investigate the role of genetic modifiers to gain a comprehensive understanding of MDs.

## ACKNOWLEDGMENTS

We thank Cécile Beust for providing the biological networks. We thank Frédérique Magdinier for critical reading of the manuscript. This study was funded by the Excellence Initiative of Aix-Marseille University—A*Midex, a French “Investissements d’Avenir” programme—Institute MarMaRa AMX-19-IET-007, The French Muscular Dystrophy Association (AFM-Telethon), and the «Priority Research Programme on Rare Diseases» of the French Investments for the Future Programme.

## AUTHOR CONTRIBUTIONS

OO, AB and MB conceptualized and designed the study. OO, SG, MC collected the dataset. OO performed the analyses. AB supervised the study. All authors contributed to the writing of the manuscript and reviewed the manuscript.

## DATA AVAILABILITY

All supplementary materials including the codes, data and results are available at https://doi.org/10.5281/zenodo.10462174.

## COMPETING INTERESTS

The authors declare no competing interests.

## Notes

https://doi.org/10.5281/zenodo.10462174

### Competing Interest Statement

The authors have declared no competing interest.

